# Human Transcription Factor and Protein Kinase Gene Fusions in Human Cancer

**DOI:** 10.1101/2020.04.09.033613

**Authors:** Kari Salokas, Rigbe G. Weldatsadik, Varjosalo Markku

**Affiliations:** Systems Pathology/Biology Research Group, Institute of Biotechnology, HiLIFE, University of Helsinki, Helsinki, Finland

## Abstract

Oncogenic gene fusions are estimated to account for up-to 20 % of cancer morbidity. Originally, oncofusions were identified in blood cancer, but recently multiple sequence-level studies of cancer genomes have established oncofusions throughout all tissue types. However, the functional implications of the identified oncofusions have often not been investigated. In this study, the identified oncofusions from a fusion detection approach (DEEPEST) were analyzed in more detail. In total, DEEPEST contains 28863 unique fusions. From sequence analysis, we found that almost 30% of them (8225) are expected to produce functional fusion proteins with features from both parent genes. Kinases and transcription factors were found to be the main gene families of the protein producing fusions. Considering their role as initiators, actors, and termination points of cellular signaling pathways, we focused our in-depth analyses on them. The domain architecture of the fusions, as well as of their expected interactors, suggests that abnormal molecular context of intact protein domains brought about by fusion events may unlock the oncogenic potential of the wild type counterparts of the fusion proteins. To understand overall effects of oncofusions on cellular signaling, we performed differential expression analysis using TCGA cancer project samples. Results indicated oncofusion-specific alterations in expression levels of individual genes, and overall lowering of the expression levels of key cellular pathways, such as signal transduction, proteolysis, microtubule cytoskeleton organization, and in particular regulation of transcription. The sum of our results suggests that kinase and transcription factor oncofusions globally deregulate cellular signaling, possibly via acquiring novel functions.

## Introduction

At any given moment, multitudes of molecular networks are activated in cells throughout the body. An important feature of these networks is that the regulation of key signaling molecules is highly concerted and deviation from homeostasis can result in diseases, such as cancer. Cancer is a complex, progressive, multi-step disorder, which stems from mutations caused by genomic instability (Hanahan and Weinberg 2011). The accumulation of genetic and epigenetic abnormalities ultimately leads to the transformation of normal cells into malignant derivatives. Two highly enriched gene groups being mutated in the majority of cancer types are protein kinases (PKs) and transcription factors (TFs) (Greenman et al. 2007; Forbes et al. 2017). PKs mediate most signal transduction events in cells by phosphorylation of specific substrates, thus modifying their activity, cellular localization, and/or association with other proteins. TFs are the “transistors” of the cellular signaling circuits, controlling the transcriptional outcome of activated signaling by binding to regulative elements of their corresponding target genes and driving or suppressing their expression. Therefore, it is easy to understand why mutational deregulation of these two gene groups can have such an impact on tumorigenesis.

In addition to harboring activating or inactivating somatic point mutations, PKs and TFs account for a large fraction of all human fusion genes involved in cancer (COSMIC, Catalogue of Somatic Mutations in Cancer, cancer.sanger.uk (Forbes et al. 2016); and dbCRID, Database of Chromosome Rearrangements in Disease (Kong et al. 2011)). Chromosomal translocations creating fusion genes are among the most common mutation class of known cancer genes, and they have long been identified as driver mutations in certain types of cancer (Futreal et al. 2004). Recently, oncogenic fusion genes (hereafter oncofusions, OFs) have been found in many hematological and solid tumors, demonstrating that translocations are a common cause of malignancy (Mitelman, Johansson, and Mertens 2004, 2007). Fusion mutations occur when two different gene regions fuse together via translocation. Examples of consequences of chromosomal fusion to protein structure range from missense mutations to expression-change inducing promoter-gene –combinations to fully functional fusion proteins with neomorphic properties. A classic example of gained functions is the breakpoint cluster region-Abelson tyrosine-protein kinase 1 (BCR-ABL1) translocation in chronic myeloid leukemia (Nowell and Hungerford 1960). Alternatively, a proto-oncogene is fused to a strong promoter, and thereby the oncogenic function is upregulated due to the strong promoter of the upstream fusion partner. This is common in lymphomas where oncogenes are juxtaposed to the promoters of the immunoglobulin genes (Vega and Medeiros 2003), and also in prostate cancer where ETS TF (ERG) is fused with TMPRSS2 regulatory sequences, thus obtaining androgen receptor (AR)-responsive expression (Tomlins et al. 2005). The current understanding favors the aberrant gene function model rather than promoter-induced over-expression.

The frequency of recurrent OFs varies depending on the specific type of cancer(Berger et al. 2011; Hammerman PS 2012; Imielinski et al. 2012; Stephens et al. 2009), but identified translocations are estimated to account for up to 20% of cancer morbidity (Mitelman, Johansson, and Mertens 2007). Recent fusion prioritization study found that in-frame transcripts were the most powerful predictor of driver fusions (Abate et al. 2014), confirming the intuition that in-frame transcripts are crucial to function. Notably, breakpoints were also observed to preferentially avoid splitting of domains. Together with frame-shift conservation, such trends could reflect a selection on fusion proteins to maintain protein stability and evade degradation pathways (Suzuki et al. 2010).

Next-generation sequencing (NGS) of genomes and transcriptomes from primary human cancer cells is constantly revealing new gene fusions that are involved in driving tumorigenesis; including examples found in colorectal carcinoma, bladder carcinoma, breast cancer and acute lymphoblastic leukemia (ALL) (Stephens et al. 2009; Muzny DM 2012; Weinstein JN 2014; Zhang et al. 2012). Furthermore, NGS has provided enough detailed sequence information of the fusion breakpoints allowing to initiate systems-level research on human oncofusions. As a result, various algorithms have been developed to mine OFs from large cancer datasets such as TCGA. However, the concordance among the different algorithms is very low that metacaller approaches utilizing consensus calls have been employed (Liu et al. 2016), which limit novel OF discoveries. Recently a new statistical method, DEEPEST(Dehghannasiri et al. 2019), was developed to overcome these limitations. In this study, oncofusions that involve PKs and TFs were selected from the data produced by DEEPEST applied to the whole TCGA dataset.

In most cases, it is not possible to draw definite conclusions about the mechanisms or extent by which any individual translocation contributes to cancer. Predicting protein function from a sequence has proven an extremely difficult task. With gene fusions, the task is even more daunting. However, an unexpectedly large number of PKs and TFs have been found to be mutationally activated or have increased expression due to gene amplification or translocation in cancer (Futreal et al. 2004). The high number of PKs and TFs with relatively low individual mutational frequency suggests either that a large number of signaling pathways can contribute to cancer, or that many PKs and TFs can regulate the same pathways when activated unphysiologically. Some additional support for this hypothesis comes from the interconnectivity of the PK-/TF-oncofusions.

In this study, fusions predicted by the AGFusion tool to produce in-frame proteins were analyzed to shed light on the protein-level implications of fusion events. The fusions were analyzed from the perspective of their domain architecture to understand likely modes of action of the novel proteins. Furthermore, known interactomes of the participating wild type proteins were used to determine possible mechanisms of action, pathways of interest, and possible treatment vectors for affecting as many different fusions as possible. As a result, multiple cellular signaling pathways were found to intersect with major subsets of these fusions, and multiple individual key interactors, such as NTRK1 with over 200 and EGFR with over 100 interacting fusions, were identified as potential targets of interest.

## Materials and Methods

### Fusion selection and annotation

Fusions that involve protein kinase genes (from Manning et al. 2012) and transcription factors (from Lambert et al) were selected from the 31,007 fusions that were identified by applying DEEPEST to the whole TCGA dataset (Dehghannasiri et al. 2019). Of these 28,862 were determined to be unique by considering Ensembl gene IDs, biotypes, chromosomal breakpoints, AGFusion assigned fusion effects, and resulting protein sequences. AGFusion was used to annotate these gene fusions to the human genome assembly GRCh38 v.89 from Ensembl. For analysis involving gene pairs, the pair entry was used in alphabetical order (e.g. ERG-TMPRSS2 instead of TMPRSS2-ERG) in all cases. Fusions were considered protein coding if both genes contributed over 30 amino acids to the product.

### Clinical Data

Clinical data for TCGA samples was obtained from the GDC data repository. The data was matched to AGFusion output data based on TCGA barcode (e.g. TCGA-WB-A80K) using a custom in-house python script. Stage information from the clinical data was simplified where possible (e.g., Stage IIA was changed to Stage II). Entries such as Stage 0, Stage X and I/II NOS were ignored. Tissue entries were simplified from detailed ICD-O 3 topographical codes to more general, e.g. C56.9 -> C56, and mapped to names accordingly. Chromosomal sequence information from GRCh38 v.89 was used to categorize breakpoints into 5 % chromosomal interval groups.

### Interactor analysis

Interactors for wild type proteins of all fusion partners were obtained from IMEx consortium (Orchard et al. 2012) and any interactions that were not confirmed to be physical by experimental methods were discarded.

Interactors were added to the interactor set from each fusion, while leaving out the fusion pair genes themselves. Annotations for interactors were obtained using Uniprot and Reactome. From Reactome, mappings to all levels of pathway hierarchy were used. Dijkstra’s algorithm (Dijkstra 1959) implemented with a custom python script was then used to establish shortest paths to Reactome root nodes for each network node. A weight of 1 was used for all network edges.

### Domain annotation

For the protein producing fusions, sequences of the protein products were produced using the AGFusion tools. Duplicate fusions based on fusion genes and protein sequence were discarded. Domains were taken from AGFusion output, and mapped to protein sequence in the wild type protein. The intactness of domains was then determined by matching the WT domain sequence to the predicted fusion protein sequence, and only full length, intact domains were picked for further study. A domain was classified as PK- or TF-specific if >= 95 % of all its occurrences were in PK or TF proteins, respectively.

### Data Visualization

Data illustrations were made with CorelDRAW, Excel, and in-house python scripts using Matplotlib and Seaborn. Cytoscape (Shannon et al. 2003) was used for creating network figures.

### Differential expression analysis

Gene expression quantification HTSeq-counts –files were downloaded from GDC data portal. Samples where OFs with intact, full-length PK, or TF domains were detected were grouped together based on fusion gene pairs. The groups were then analyzed with DESeq2 (Love, Huber, and Anders 2014) using other samples with protein producing non-PK/-TF fusions as controls, and the resulting significantly differentially expressed genes (FDR corrected p <= 0.01) were annotated with GO terms and Reactome pathway identifiers. Z-score value for pathway level over-/underexpression was calculated by a method used in GOplot (Walter, Sanchez-Cabo, and Ricote 2015) i.e.by deducting the number of underexpressed genes from the number of overexpressed genes and dividing the result by the square root of the number of significantly changed genes (FDR corrected p <= 0.01).

## Results

### Detection of oncofusions from TCGA dataset reveals enrichment of PK and TF fusions

In this study, we focused on protein producing OF genes. Translocation of two chromosomal regions can result in either in-frame or out-of-frame OFs (Fig1A). To characterize the proteins produced by currently known OFs in the TCGA dataset, which currently contains data from 33 different cancer projects, we embarked on an analysis to understand the potential functional space of the protein producing fusions (Fig 1B), and especially those that involve either a PK or a TF, or both (PK-TF fusions).

**Figure 1:**
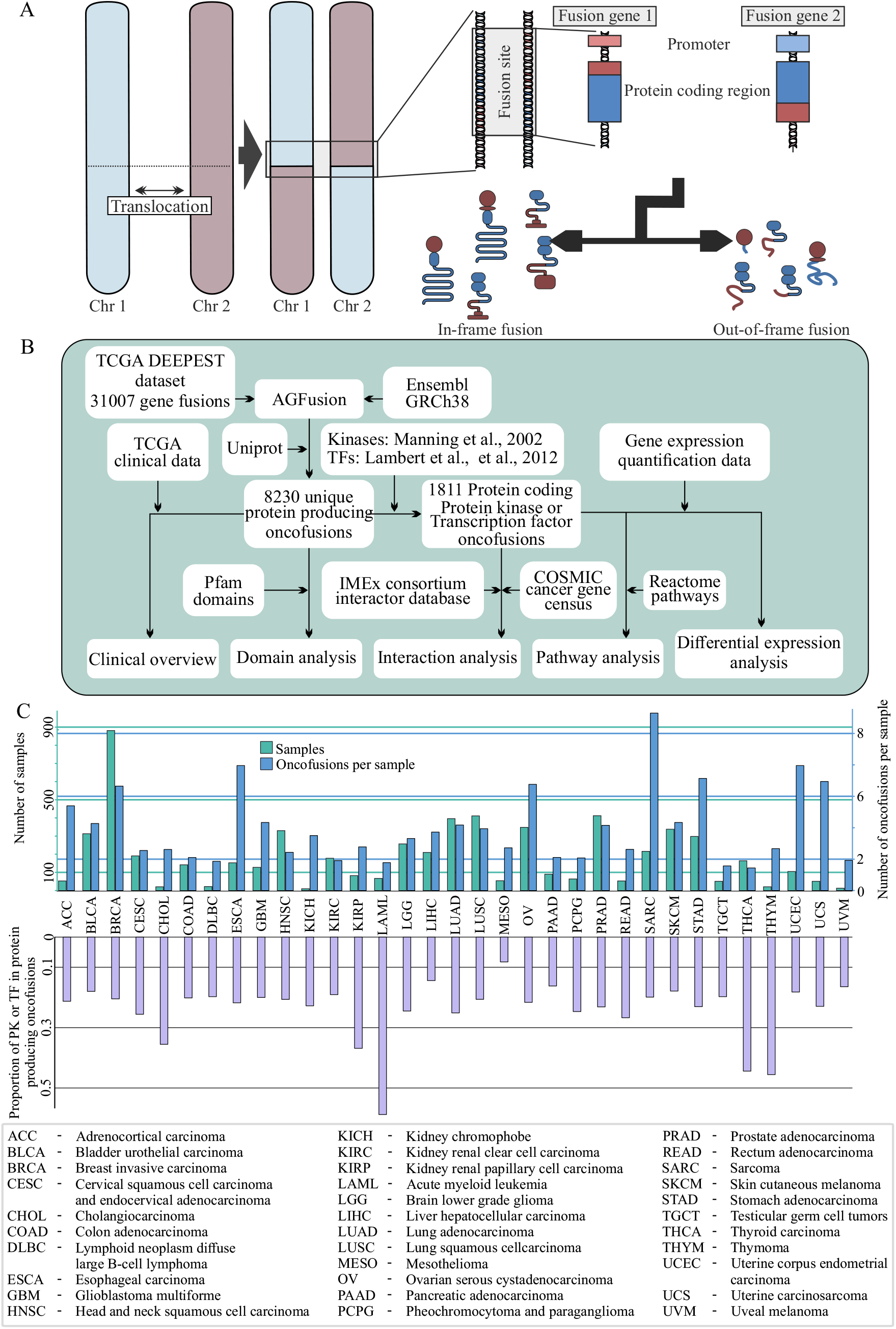
Schematic illustration of the gene fusions, workflow, and the number of gene fusions in human cancer. **A)** Schematic description of gene fusions formation. Fusions are formed mainly via balanced and unbalanced chromosomal rearrangements, such as translocations, deletions, inversions and insertions. This usually leads to formation of a fusion gene with the 5’ end of Gene 1 and 3’ end of Gene 2. If the fusion occurs between two protein coding genes, depending on whether the reading frame is violated, and where exactly the fusion occurs, a fusion protein may be transcribed with features and domains from both partners. Other possible outcomes include full or truncated 3’ gene under the control of the promoter of the 5’ gene. **B)** Workflow used in this study. Analysis progressed from the total set of gene fusions discovered by the DEEPEST method and moved towards more specific kinase / TF containing, protein producing oncofusions. We started with TCGA data-based fusion set from Dehghannasiri et al. (2019), for which we generated protein sequences with AGFusion. Domains were added by matching sequence to Uniprot proteins annotated with Pfam domains, after which non-unique entries were dropped. In total, the dataset included 8,230 unique protein producing fusions. Fusions were classified as protein producing, if both gene fragments were predicted to produce > 30 AAof protein sequence. From this set, the two most prominent protein groups were protein kinase and transcription factor, and thus we focused further analysis on the 1,811 unique protein kinase or transcription factor containing fusions, using the full protein producing fusion set for comparison. Known interactions for wild type fusion proteins were obtained from IMEx consortium, and used for estimating maximal foreseeable effect on signaling pathways from Reactome. Finally, TCGA gene expression quantification data was used to probe possible observable effects of kinase/TF fusions, using other protein producing fusions as background. **C) Top:** Breakdown of samples and fusion mutations by TCGA project. Largest single contributor of samples with fusions was TCGA breast invasive carcinoma project (BRCA), which had the highest number of samples and identified fusion mutations. **Bottom:** Proportion of protein producing fusions that include PK or TF genes. Protein producing is defined as both 5’ and 3’ fusion partner fragments producing over 30 amino acids of their respective wild type proteins.

The DEEPEST dataset included 31,007 fusions detected from 6,123 cancer samples. Of these, 28,862 were unique fusions (Fig 1C, upper panel). Among the unique OFs, 29 % (8,230) were predicted to not only retain frame, but to also produce potentially functional proteins, where both genes contributed over 30 in-frame amino acids (Fig 1B, Supplementary table 1). The limit of 30 amino acids was the length of the shortest non-repeat domain present in any of the fused proteins. Examining the resulting protein producing OF set, we noticed an abundance of those involving PK or TF. Indeed, these fusions constituted 1,811 protein producing OFs (Fig 1C, lower panel). Generally the proportion of was under 0.3, except in the PK/TF –fusion prone cancers acute myeloid leukemia, cholangiocarcinoma, thyroid carcinoma, and thymoma. The number of OFs per sample varied across cancer types. The types most prone to protein producing fusions were sarcoma (SARC) with an average of 3.7 protein producing fusions per sample, esophageal carcinoma (ESCA: 3.5 fusions), uterine corpus endometrial carcinoma (UCEC: 2.9), stomach adenocarcinoma (STAD: 2.8), breast invasive carcinoma (BRCA: 2.7), uterine carcinosarcoma (UCS: 2.6), and ovarian serous cystadenocarcinoma (OV: 2.5).

Due to the prevalence of PK and TF genes in the fusions, we next investigated if they are enriched in particular cancers. While in most cancers PK/TF fusions made up around 20-25 % of all protein producing OFs, the percentage reached 60 % in acute myeloid leukemia (LAML) samples, 46 % in thymoma (THYM), 45 % in thyroid carcinoma (THCA), and 37 % and 36 % in kidney renal papillary cell carcinoma (KIRP) and cholangiocarcinoma (CHOL) respectively (Fig 1C). Acute myeloid leukemia is well known as an OF-prone cancer Gao et al. (2018). However, aside from the four fusions detected between ABL1 and BCR, the high percentage was mostly TF-driven, with KMT2A, RUNX1, and RARA being found in 9, 6, and 4 fusions respectively. This is in contrast to the peak in THCA, which is driven by 12 BRAF fusions, 11 fusions of RET, 6 of NTRK1, and 5 of NTRK3, among 8 other protein kinases.

### Reading frame retention is common in PK and TF oncofusions

The 31,007 fusions consisted of 23,354 unique gene pairs and 14,632 individual genes; 14,338 of the pairs did not have any protein producing fusions. The top protein producing fusion was RPS6KB1-VMP1 with 13 unique protein producing fusions in the dataset, all the others having less than 10 (Fig 2A). There were 47 fusion gene pairs that were predicted to produce protein in at least 4 fusions, 159 in 3 fusions, 835 in 2, and 7975 in 1 fusion. Out of the 32 fusion gene pairs that produced 4 or more unique proteins, 15 were PK/TF fusions.

**Figure 2:**
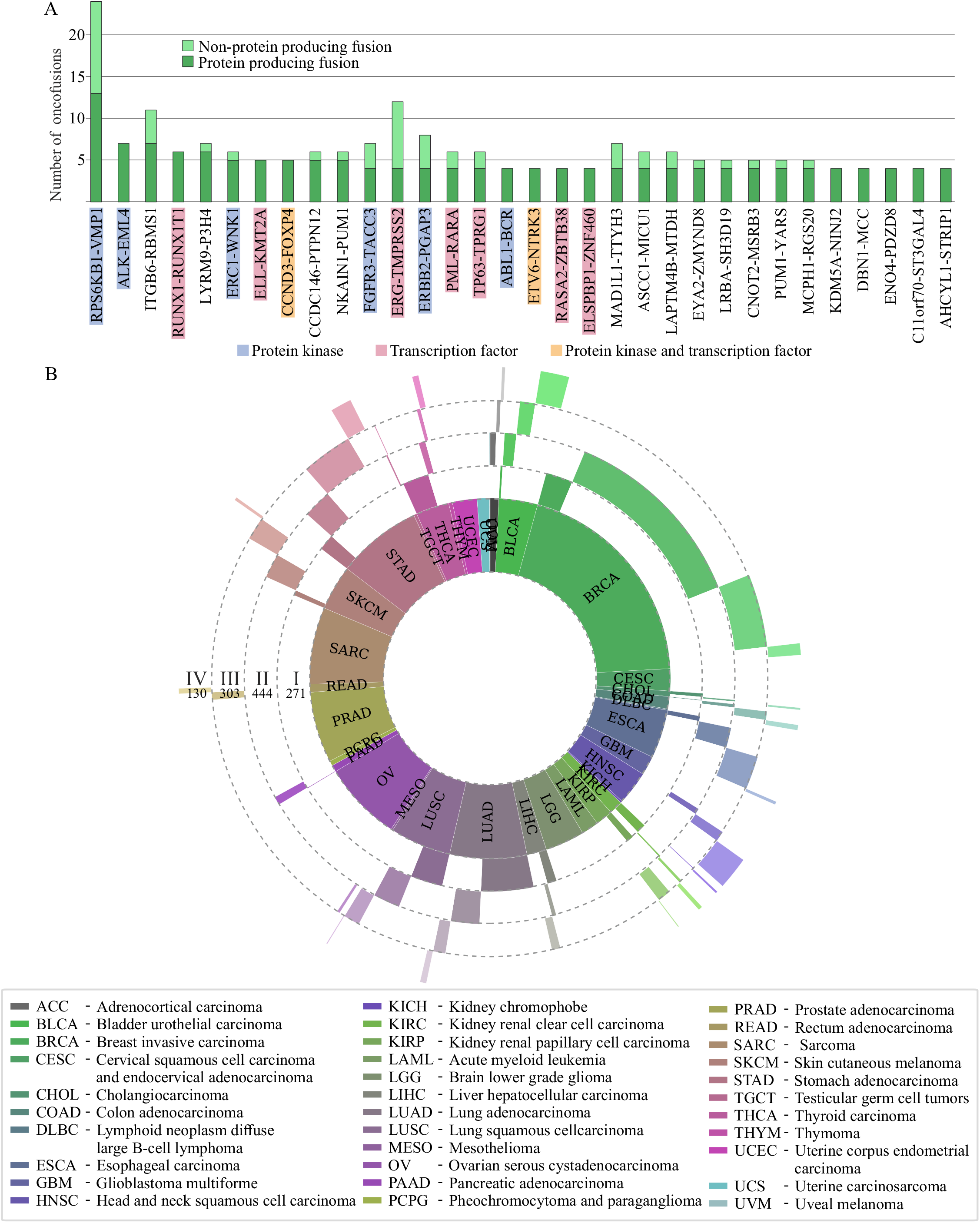
Clinical characterization of protein producing oncofusions by cancer stage. **A)** The most common protein producing gene pairs in oncofusions. In total, protein producing fusions were comprised of 23,354 unique gene pairs predicted to produce one or more unique protein products. The most common pair was RPS6KB1-VMP1, with over 10 unique proteins, followed by ITGB6-RBMS2 and ALK-EML4 with 7 each, and LYRM9-P3H4 and RUNX1-RUNX1T1 at 5. Kinase and TF fusions were common in top protein producing gene pairs, illustrated by blue shading for the presence of a protein kinase in gene pair, red for TF, and orange for both. **B)** Sunburst diagram of project and stage distribution of PK/TF oncofusions. The innermost layer represents the number of fusions in each project. The layers radiating out are the proportion of fusions detected in Stage I, II, III, and IV samples, in order from in to out. Total numbers of fusions from each stage is marked under the stage indicators.

To better understand the behavior of prolific gene pairs, we next mapped tissue annotations from TCGA to fusions of each gene pair based on barcodes from samples, where a fusion of the gene pair was present. In contrast to RPS6KB1-VMP1 and ITGB6-RBMS1, which were seen in samples from 6 different cancers, 7,055 pairs were seen in samples of only one cancer type. Out of these cancer-specific fusions, 38 were predicted to produce 2 or more unique proteins (with ERG-TMPRSS2 predicted to produce 4 different unique proteins, supplementary table 2). PK/TF fusions featured 1449 different PK or TF genes, ERG being the most common TF, and ERRB2 the most common PK (supplementary table 3). Between 84 and 97 percent of oncofusions in each cancer project were unique, highest being sarcoma with 97 % unique gene pair combinations, and thyroid carcinoma the lowest with 84 % (supplementary table 4). Protein producing fusions followed a similar theme, unique protein producing fusions making up between 23 and 52 % of all oncofusions in each given cancer project (supplementary table 4).

We next looked in more detail what cancer stages PK and TF fusions were detected in. The most prominent group was stage II breast invasive carcinoma, which also had the most samples in the data set (Figure 2 B). In total, of the 1,811 PK and TF fusions, 271 were found in stage I samples, 444 in stage II, 303 in stage III, and 130 in stage IV. On average, samples had 0.30 PK/TF fusions per sample. However, in some cancers, PK or TF fusions are enriched towards the more severe stages. Discounting stage groups with less than 10 samples, 4 groups had more than 0.6 fusions per sample. In particular, ESCA stage III samples in particular had 0.76 PK/TF fusions per sample, while STAD and BRCA stage IV samples had 0.69 and 0.65 respectively, and STAD stage I had 0.61 (Supplementary table 5). The distribution of protein producing OFs mirrored that of PK/TF fusions quite closely (supplementary figure 1A). In terms of chromosomal breakpoint locations, those in the PK/TF fusions varied compared to all protein producing fusion mutations, but the prominent role of PK/TF fusions is illustrated by overlapping hotspots (supplementary figure 2).

### Intact, in-frame domains are commonly retained in OFs

To understand the contribution of each OF to the overall development or survival of cancerous cells, the functional consequences of any given mutation and its impact on the pathways the proteins are involved in must be understood. To this end, we analyzed all identified unique protein producing fusions, and the full-length, in-frame domains of the fusion proteins.

While AGFusion does predict protein sequence for each fusion partner, and corresponding conserved or lost domains, a domain is counted as conserved already if only 5 amino acids are included in the sequence. To adapt this to the study of full-length domains, we first mapped the Pfam identifiers of the domains to sequences in the wild type proteins from Uniprot. The domains were then defined as conserved only if the full sequence was present in the fusion protein. This resulted in 10,100 conserved domains in all protein producing fusions. Over 50 % (5,373) of these domains are in PK/TF fusions, which account for 22 % of all protein producing fusions (supplementary tables 1, 6), suggesting overall domain count strongly favors PK and TF genes, perhaps indicating that these fusions produce more functional proteins in comparison to all protein producing fusions.

The most conserved domain was the protein tyrosine kinase domain (Fig 3A, supplementary table 6), which was conserved in 159 fusions. This was followed by the PH domain, a common domain in intracellular signaling proteins and proteins of the cytoskeleton, and the protein kinase domain. To assess retention of non-obvious PK or TF domains, we classified domains as PK or TF specific if over 95% of the copies were found in PK or TF halves of the fusion proteins. This resulted in 622 copies of 131 different TF-specific domains predicted to exist in the fusions, compared to 455 copies of 44 PK-specific domains. Most common TF domains were zinc finger C2H2 type, KRAB, and HLH DNA binding domains, present in 59, 45, and 43 copies respectively. Many TF domains, such as KRAB, are involved in both transcriptional activation and repression, depending on the molecular context.

**Figure 3:**
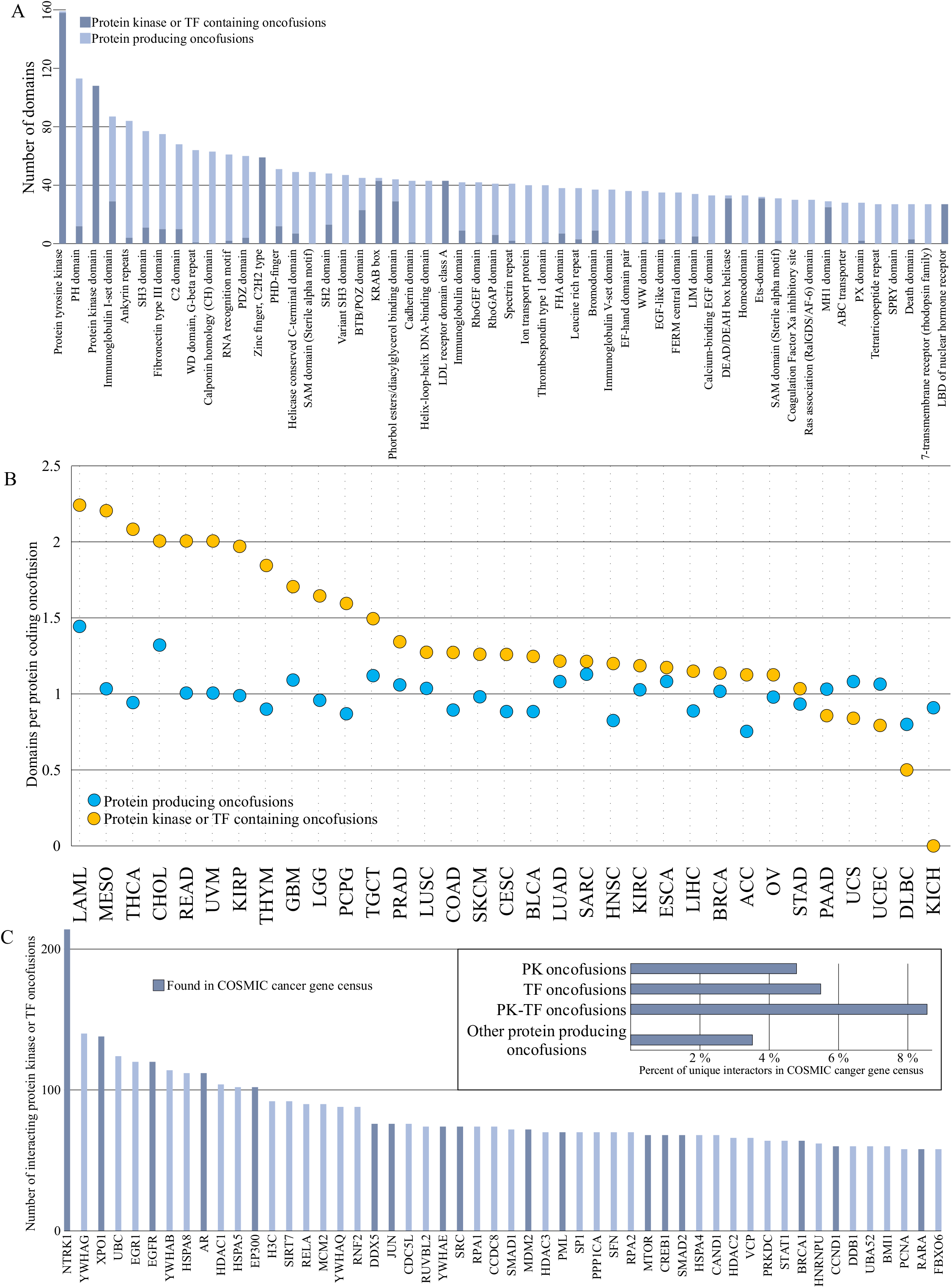
Domain analysis of protein producing fusions. **A)** Intact, full-length domains identified in unique protein producing fusions. In total, 10,100 intact domains were detected. The protein tyrosine kinase domain was the most prevalent with 159 identifications. In addition, protein kinase domain was detected with 108 copies each. Kinase or TF specific domains included 44 and 131 unique domains, respectively. 455 copies of kinase-specific domains were seen, and 622 of TF specific. Kinase domains focused more on the two top kinase domains, whereas TF domains were a much more evenly distributed group, the top TF-specific domain, C2H2 type zinc finger, having 59 copies. **B)** The number of domains per protein coding oncofusion in the TCGA projects. **C)** Most common interactors of protein kinase/TF fusions. Y-axis describes numbers of unique protein producing fusions, where one or both of the fusion partner WT genes interact with the protein. Top right inset: Proportion of interactors found in COSMIC cancer gene census is higher in both protein kinase and transcription factor fusions, and most common in fusions between protein kinase and transcription factor genes.

On average, protein producing fusions in samples of most cancer projects tended to have close to 1 intact, full length domains per protein producing OF. PK/TF fusions on average had more intact domains in all except for 5 projects (Fig. 3B). On average, fusions in all projects tended to have between 1 and 2 intact domains, while PK/TF fusions featured a slightly larger average. Although some cancers do appear to have particularly many domains, this is mostly due to low count of fusions detected in the project. Exception seems to be acute myeloid leukemia, with 47 detected protein producing fusions, 28 of which contain either a PK or a TF. Most striking differences being seen in mesothelioma, thyroid carcinoma, rectum adenocarcinoma, and uveal melanoma with 1.17, 1.14, 1.0, and 1.0 more retained domains on average in PK/TF fusions compared to protein producing fusions, respectively.

On the cancer project level, thyroid carcinoma had the highest percentage of PK domains (19 % of all domains identified in the 97 samples of the project, supplementary table 7, supplementary Figure 3), which totaled to 34, only exceeded by breast invasive carcinoma with 92 PK specific domains (4 % of all BRCA domains), and lung adenocarcinoma (LUAD) with 35 (6 %). Proportion of TF domains varied less. Kidney renal papillary cell carcinoma had 15 % of its intact domains in the TF-specific set, followed by acute myeloid leukemia with 12 %, and rectum adenocarcinoma and prostate adenocarcinoma, both at 11 %. Aside from prostate adenocarcinoma, these projects had < 50 samples in the TCGA dataset.

### Interactors of fusion partners can point to impact of OFs

To understand what kind of implications the functional changes of lost / conserved PK or TF specific domains in new combinations could have for the cell and the organism as a whole, we next analysed the interaction networks of the wild type proteins in PK/TF fusion set.

Although PK or TF fusion proteins are likely to lose domains necessary for these interactions to form, they are also likely to instead gain domains facilitating new interactions. We took the known, experimentally validated interactomes of the wild type proteins from the IMEx consortium (Orchard et al. 2012). We treated the resulting interactor set as the hypothetical maximal foreseeable effect set, which consisted of interaction partners that may have an effect on the fusion protein, or that the fusion protein may have an effect on.

From this set, we found interactors that were particularly prominent. PKs, such as NTRK1, EGFR, and SRC, as well as various TFs, like NFKB3 and SP1 were among the top results (Fig 3C, supplementary table 8). The list is mostly made up of other kinases or transcription factors, with NTRK1 potentially interacting with over 200 individual, unique fusions. Genes found in COSMIC cancer gene census were more common towards high numbers of potentially interacting fusions. Through these interactions it is possible to identify significant central nodes through which multiple different fusions in different cancers may affect the growth of the tumor. For example, the second most common possible interactor, YWHAG, is a common regulator of signaling pathways. Approximately 7.7 % of all interactors of protein producing fusions were found in COSMIC cancer gene census, whereas the percentage rises to 29 %, and to 40 % if we consider only the 100 and 10 most common protein producing fusions respectively. Interactors of PK or TF fusions were more often seen in the cancer gene census, than those of other protein producing fusions (Figure 3C upper right inset).

### Pathway analysis of OF interactors highlights signal transduction and regulatory functions

Next, we combined Reactome pathway data to the interactor set, and built hierarchic networks of the found pathways (Fig 4A). Considering the dataset, we focused on one network root node: signal transduction, and its descendants up to 7 links away. Another root node, gene transcription, can already be seen on this scale, which is unsurprising considering the inclusion of many TF fusions, and the interplay of signal transduction and gene transcription. For each Reactome pathway, we calculated an interactor count by adding together the number of potentially interacting fusions for each protein in the pathway.

**Figure 4:**
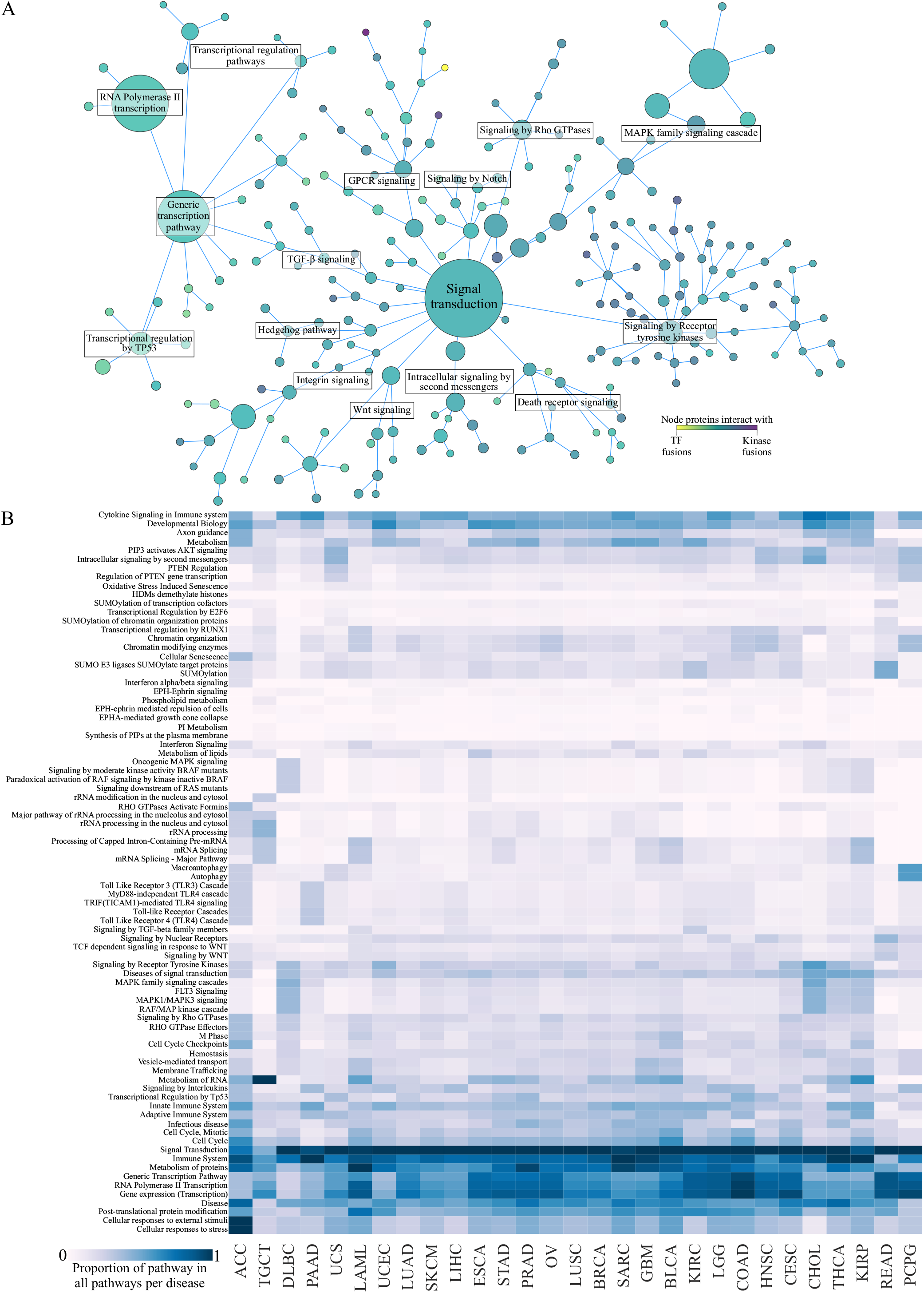
Functional potential of Kinase/TF fusion interactors. **A)** Interactors mapped to Reactome pathways. The interactors produced hits in almost 2000 pathways. Most prominent hits were centered around signal transduction pathways, which links to transcription events via TGF-β signaling pathway. The size of the node is directly proportional to number of fusions with interactors identified with the annotation from Reactome database. The used annotation file contained annotations for all levels of Reactome hierarchy. Included in the figure are pathways up to 7 steps away from the signal transduction root node. The node size is directly proportional to the sum of oncofusion interactors, and the count of fusions that interact with them. **B)** Relative frequency of each pathway per TCGA project on a scale from 0 to 1 (1 being the pathway with most interactors). While pathways with the most potential interactors of fusions identified are the same in majority of the projects, different subpathways are seen in different projects, such as oncogenic MAPK signaling in DLBC and KIRP, or PI3-Akt signaling in UCS and CHOL.

Particularly enriched were proteins related to signal transduction, where interactors were detected in 15 branches from the root. Especially prominent pathways are those relating to receptor tyrosine kinase signaling with potential interactors from 1,230 unique PK/TF oncofusions), PI3K-Akt (1,220), Rho GTPases (1,095), Integrin (1,156) and GPCR (905) signaling, as well as MAPK family signaling cascades (1,053). Multiple smaller, but significant pathways such as Hedgehog, Notch and Wnt pathways are also seen. Generic transcription pathways and their related pathways, such as transcriptional regulation and RNA polymerase II transcription, are very prominent as well, with 1,503 PK/TF oncofusions.

The proportion of interactors from each pathway varied slightly between different cancers (Fig. 4B). While signal transduction was the most common pathway in most cancers, different signaling cascades, such as MAPK cascades or TLR cascades featured much more variation, pointing to relative enrichment of different pathways in different fusions, and perhaps to cancer-specific effects of unique gene-pair mutations in said cancers.

### Oncofusions lead to distinct changes in gene expression

To understand if intact PK or TF domains had a recognizable and distinct downstream effect on gene expression, differential expression analysis was performed. Gene expression quantification result files were downloaded from GDC data portal, and divided into groups based on PK/TF fusion gene pairs. Only gene pairs detected in at least three different samples with results available were used.

In order to discover variation specific to PK/TF fusion gene pairs, samples were divided into those where PK/TF fusions were found in, and those with other protein producing OFs. From the latter 100 were randomly picked as a background set. Non-primary tumor samples were also discarded. This resulted in six gene pairs: RPS6KB1-VMP1, MERTK-TMEM878, FGFR3-TACC3, ERC1-WNK1, and BRAF-SND1. The analysis was repeated three times for each gene pair with new randomized background set every time. For comparison, a parallel differential expression analysis with any PK and any TF fusion was performed.

Results were filtered based on q-value < 0.01, and changes measured in log2 fold change. As expected, the PK and TF groups produced only small changes in expression patterns, presumably due to differences being averaged out in a large, heterogeneous sample group. In comparison, gene pair groups of interest produced visible effects on genetic level (supplementary table 9).

To further eliminate small-scale changes, any gene with log2 fold change between −3 and 3 in all samples was discarded from the data. As a result, several clusters of genes with positive or negative log2 fold changes in all of the three analyses of each gene pair can be seen (Fig. 5A). In total, 199 genes were differentially expressed in one or more of the gene pair groups, of which 4 were found in COSMIC cancer gene census, and 28 were associated with cancer in DisGeNet. Three of the genes were found in both (supplementary table 10). Out of the six gene pair groups, MERTK-TMEM878 appears most similar to the protein producing fusion containing sample controls. In general, only a few large differences were seen, which suggests tumor mechanisms in the samples characterized by these fusion pairs were not dramatically different from all other tumor samples, but each introduces their own variation on the theme. Next, fold changes were averaged over the three samples of each group and genes mapped to GO terms via Ensembl Biomart. Sum effect for each GO term was calculated by assigning −1 to each log2 fold change in filtered genes below zero, and +1 for above. MERTK-TMEM878 and ERC1-WNK1 did not feature many changes on GO level (supplementary table 11), and were discarded from further study. Out of the six gene pairs studied, RPS6KB1-VMP1 was seen most widely, in 7 cancer projects (BLCA, BRCA, CESC, HNSC, LUAD, LUSC, STAD), FGFR3-TACC3 and ERC1-WNK1 in 5 (BLCA, CESC, HNSC, LIHC and LUSC; and ESCA, HNSC, LUAD, LUSC and UCEC, respectively), while MERTK-TMEM87B and ERC1-RET were seen in 2 (CESC and OV; BRCA and THCA, respectively). BRAF-SND1 was only seen in THCA samples. The results point to pan-cancer modes of actions for the most widely seen, while the effects of BRAF-SND1 may be more tissue-specific.

**Figure 5:**
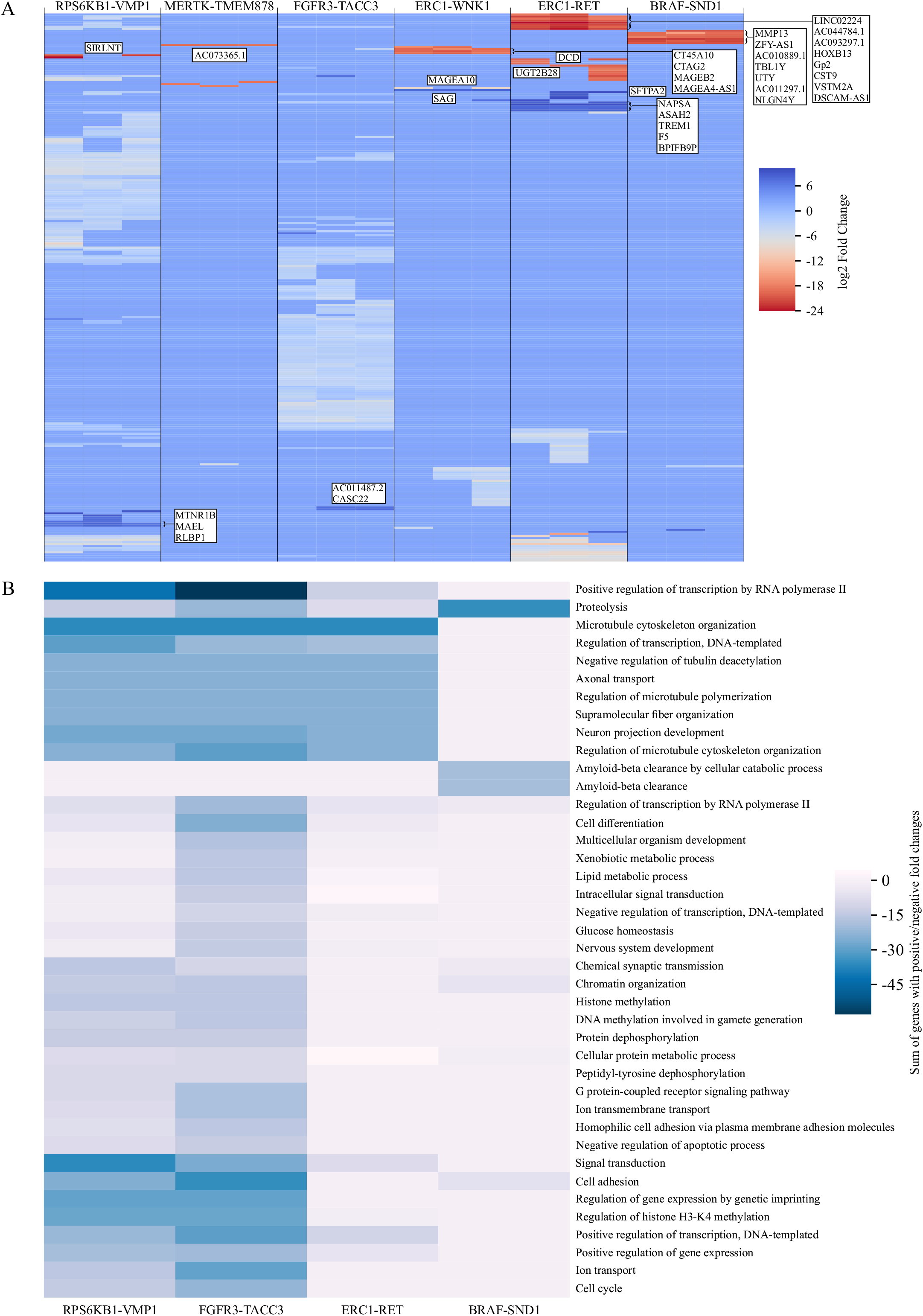
Results of differential gene expression analysis. **A)** Overall figure of the differentially expressed genes in three replicates of each gene pair group. Gene pair groups with the most positive and negative fold changes are indicated. Data was filtered with a q-value threshold < 0.01, and absolute log2 fold change of over 3. Filtered out entries were assigned 0 as value to highlight bigger fold changes. **B)** GOBP analysis of the differentially expressed genes. Significantly, differentially expressed genes were counted as either −1 or +1 for each GO term associated with them, and results summed.

RPS6KB1-VMP1, FGFR3-TACC3, ERC1-RET, and BRAF-SND1 however each produced a similar, yet unique profile (Fig. 5B, supplementary table 11). Samples of the gene pairs were mainly characterized by lack of regulation. E.g. transcription regulation and dephosphorylation are both lower than in other protein producing tumor samples. In particular, functions related to regulation of gene expression, cell adhesion, and signaling, all relevant to tumor development and survival, were downregulated. Taken together, the data presented here is an indication of the potential of gene fusions, as well as a hint at possible impacts.

## Discussion

We examined the gene fusion landscape in human cancer (TCGA datasets). Gene fusions are among the most common mutation classes of known cancer genes (Futreal et al. 2004), found both in hematological and solid tumors. Although the fusions can drive cancer via expression level changes when an oncogene is fused with a strong promoter such as TMPRSS2-ERG fusions in prostate cancer (Guo, Gui, and Cai 2011), we find that the majority 19,911 of the 28,863 oncofusions are in-frame mutations between exonic regions of two protein coding genes. In total, over 9,000 gene pairs were seen participating in fusions that were predicted to produce intact, potentially functional proteins. TCGA solid tumor samples tended to have fusions producing in-frame proteins of adequate length approximately 20 % of the time in all diseases (Figure 1C). Equal distribution across stages may hint at protein producing fusions being early events in the development of the tumors from which they were identified. We identified several particularly prolific fusion gene pairs, among them capturing also several of which have been featured prominently in literature. Most prolific protein producing OF gene pairs were found to feature either a protein kinase or a transcription factor (Fig. 2C), further validating the previously suggested idea that protein kinase and transcription factor fusions constitute to a major fraction of the oncofusions.

We next moved on to characterize the structure of produced fusion proteins, to understand the protein-level consequences of the mutations, and thus the possible impact on protein activity in a cellular context. Particularly abundant protein groups among all protein producing fusions were PKs and TFs, which has been noted in previous studies as well (Gao, 2018). We therefore decided to focus on their fusions in particular.

Intact, full-length domains were abundant in the predicted fusion proteins. Especially protein kinase domain was very prominent, featured in 159 unique protein producing fusions (Figure 3, supplementary table 6). As the kinase domain is usually in the C terminal of the protein, fusion mutations can easily cause kinase domains to be mislocalized due to localization signals from the fusion partner protein, or deleted membrane-spanning regions of the original kinase, for example. In addition to the protein tyrosine kinase domain, 43 other PK-specific domains were identified, bringing the total number of PK-specific domains to 455. In comparison, TF-specific domains consisted of a wider variety of individual domains, with 622 copies of 131 different domains.

Although fusion proteins are likely to lose domains that facilitate the validated interactions of the wild-type proteins, this is not necessarily the case. Receptor tyrosine kinases for example are commonly at the 3’ end of the new fusion gene, and the breakpoint often occurs just on the 3’ side of the region coding for transmembrane part of the receptor. This could cause the kinase domain and intracellular protein-protein interaction domains to end up in a localization dictated by the 5’ gene. This, in turn, would lead to activation in an inappropriate place and/or at an inappropriate time. Similarly affected may be proteins shuttling between nucleus and cytoplasm as a response to an outside signal: they may end up perpetually trapped in the cytoplasm or the nucleus, or shuttled between the two in atypical conditions. To understand what kind of impact protein domains in novel environments might have, we next looked at already known interactors of all fusions (Figure 3C, 4). By grouping together interactors of both wild type proteins of each fusion, we were able to estimate the maximal set of currently foreseeable interactors of the fusion protein. I.e. phosphorylation targets, complex components etc. We found genes mentioned in COSMIC cancer gene census to be enriched in both PK and TF fusions when comparing to other protein producing fusions, and occurring at a much higher rate in the interactor set of fusions between PK and TF genes. Same trend is reflected in CGC genes being more common the higher the number of potentially interacting fusions is. Possible roles of the interactors were then investigated via the Reactome pathway database. Interactors of PK and TF fusions were heavily concentrated around signal transduction pathways (Figure 4).

To gain insight into whether the deductions so far were valid, we next used TCGA transcription quantification data to dig into the effects of specific gene-pair fusions with intact kinase or transcription factor specific domains. We discovered observable changes in gene expression, when comparing fusion groups against other protein producing fusions (Figure 5), and effects that were seen to produce a noticeable impact on a GO-term level as well, pointing to lower activity of various regulatory functions with many of the domain-containing gene pair oncofusions.

The conservation of domains may suggest conserved active functions, such as those of the kinase domain, potentially linked to inappropriate dimerization domains, target recognition domains, or domains that alter the entire molecular context of the novel fusion protein by targeting it to the wrong cellular compartment, membrane, or membrane raft. Through analyzing interactors of wild type proteins, we have identified multiple common interactors. If these interactions rely on the intact domains of the fusion protein, we can assume they represent possible pan-fusion drug targets, with which a multi-cancer effect may be achieved.

Taken together, we have now created and characterized the largest dataset of kinase and transcription factor oncofusions. This database will work as the foundation for molecular cloning and characterization of the PK- and TF-oncofusions using biochemistry, proteomics and cell biology – and a baseline hypothesis for the expected results.

---

**Supplementary figure 1.**
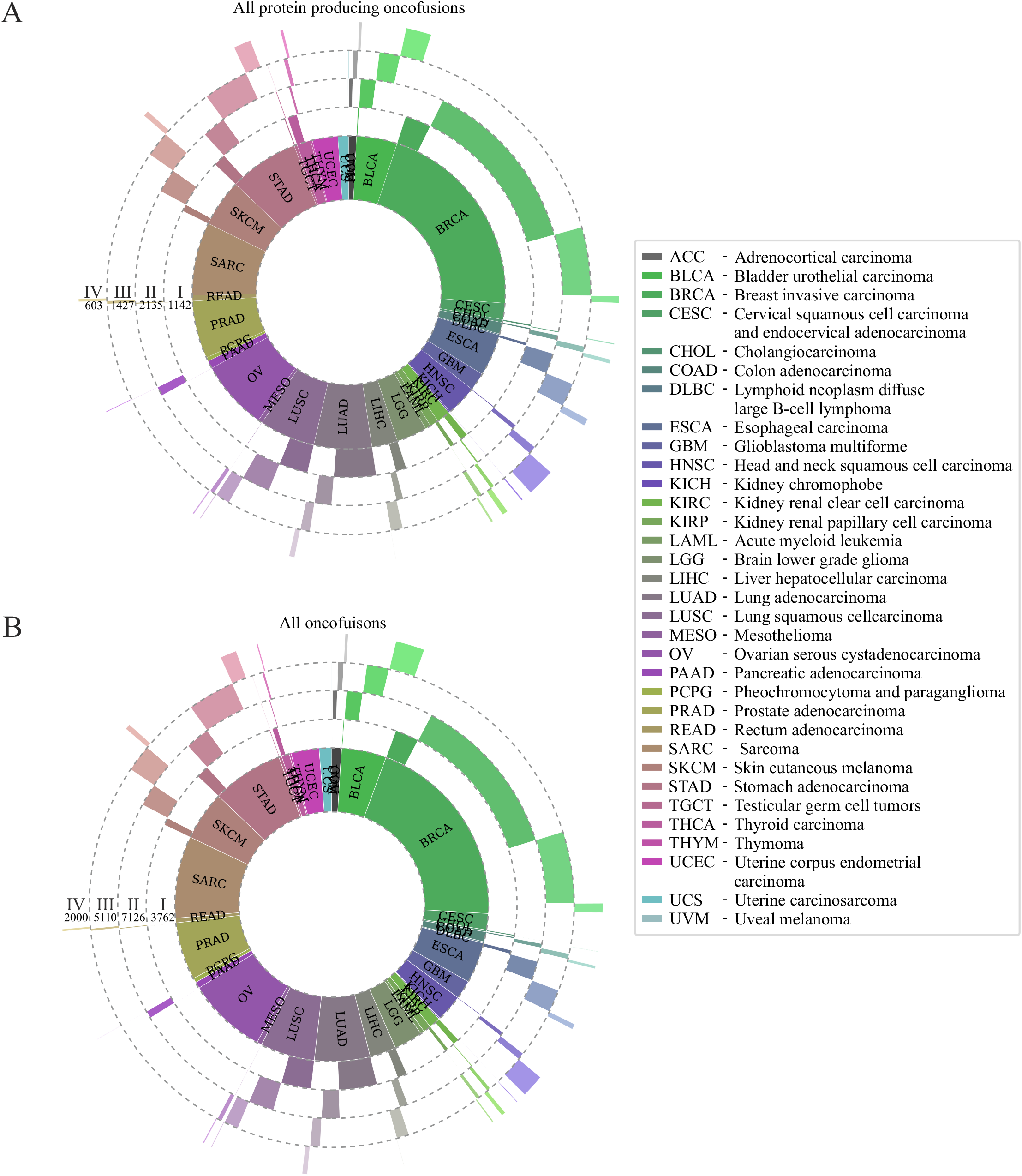
**A)** Stage and project distribution of all oncofusions found in the TCGA dataset. **B)** Distribution of protein producing oncofusions across TCGAcancer projects. Distribution seen is nearly identical to that seen with all fusions, and similar to that of PK/TF fusions (figure 2A).

**Supplementary figure 2:**
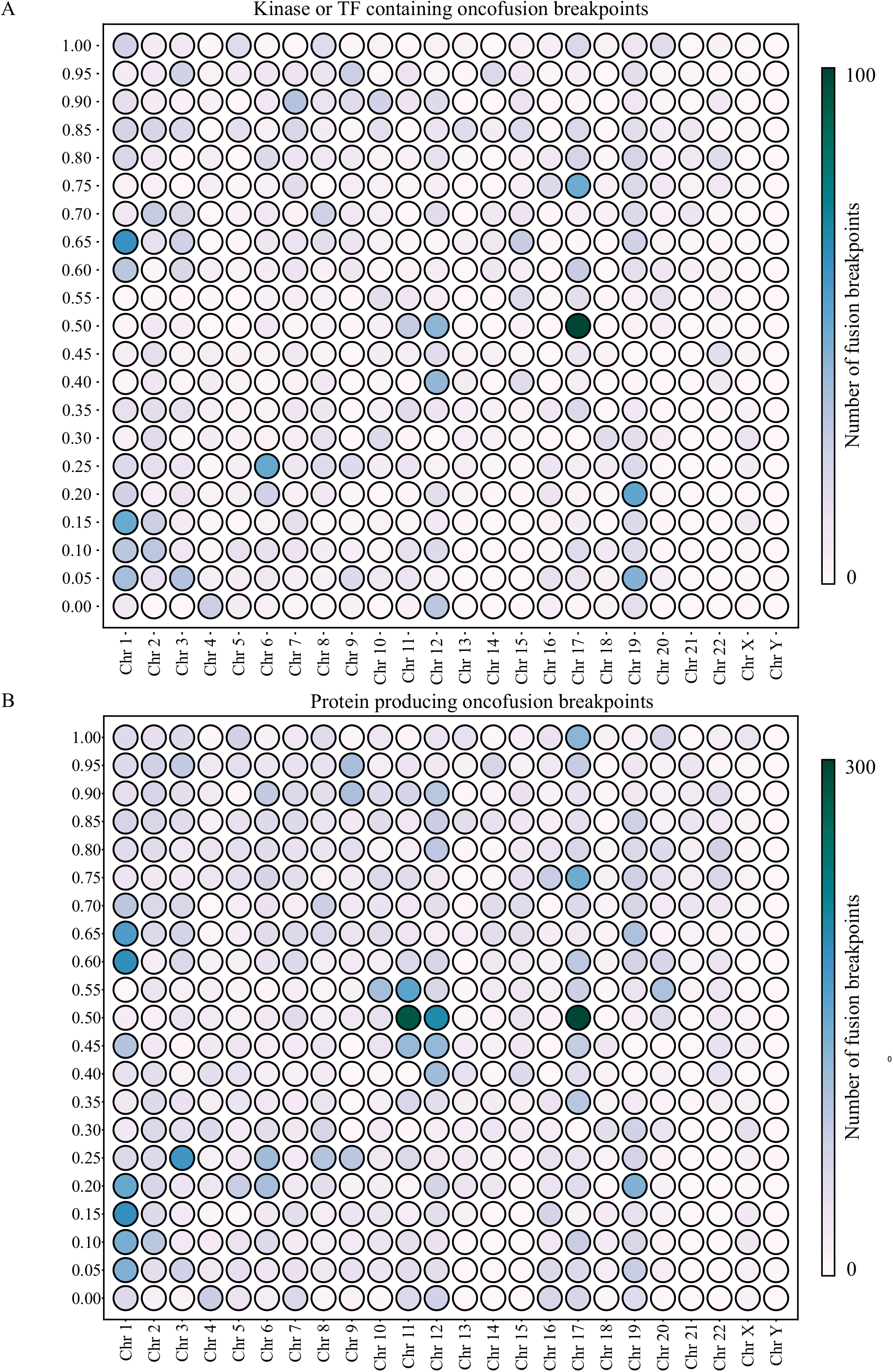
Chromosomal distributions of fusion breakpoints as percentage of chromosome length. **A)** PK/TF fusions feature one very prominent hotspot around the 50 % mark of chromosome 17, and several less intense ones. The 50 % spot on chromosome 17 is driven mainly by oncofusions with either ERBB2 or RARA as one fusion gene. **B)** Protein producing fusion breakpoints feature prominent hotspots in common with PK/TF fusions, and in addition several unique spots, where not many PK/TF fusions are present.

**Supplementary figure 3.**
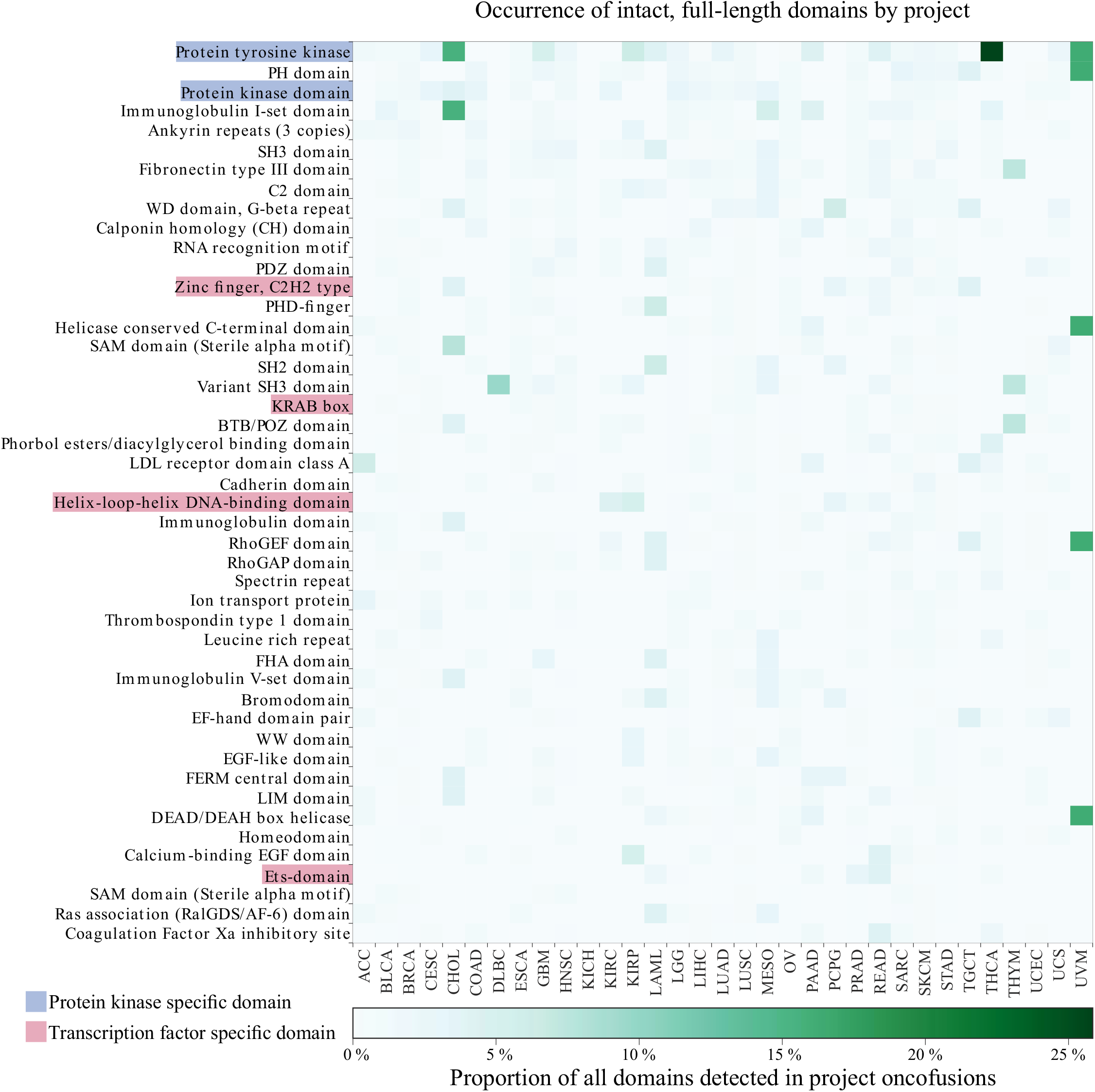
Distribution of intact, full-length domains in fusion proteins per TCGA project. Counts are minmax normalized to 1 per project to account for variable number of fusions in each project.

